# Sequence learning is driven by improvements in motor planning

**DOI:** 10.1101/523423

**Authors:** Giacomo Ariani, Jörn Diedrichsen

## Abstract

The ability to perform complex sequences of movements quickly and accurately is critical for many motor skills. While training improves performance in a large variety of motor-sequence tasks, the precise mechanisms behind such improvements are poorly understood. Here we investigated the contribution of single-action selection, sequence pre-planning, online planning, and motor execution to performance in a discrete sequence production (DSP) task. Five visually-presented numbers cued a sequence of five finger presses, which had to be executed as quickly and accurately as possible. To study how sequence planning influenced sequence production, we manipulated the amount of time that participants were given to prepare each sequence by using a forced-response paradigm. Over 4 days, participants were trained on 10 sequences and tested on 80 novel sequences. Our results revealed that participants became faster in selecting individual finger presses. They also preplanned 3-4 sequence items into the future, and the speed of pre-planning improved for trained, but not for untrained, sequences. Because pre-planning capacity remained limited, the remaining sequence elements had to be planned online during sequence execution, a process that also improved with sequence-specific training. Overall, our results support the view that motor sequence learning effects are best characterized by improvements in planning processes that occur both before and concurrently with motor execution.

**New & Noteworthy:** Complex skills often require the production of sequential movements. While practice improves performance, it remains unclear how these improvements are achieved. Our findings show that learning effects in a sequence production task can be attributed to an enhanced ability to plan upcoming movements. These results shed new light on planning processes in the context of movement sequences, and have important implications for our understanding of the neural mechanisms that underlie skill acquisition.

## Introduction

Many everyday skills require the production of sequences of individual movements. For example, typing on a computer keyboard consists of producing serially ordered sequences of finger presses. While novices tend to type slowly, taking time to select every single press and frequently going back to correct their mistakes, experts can type quickly and flawlessly, executing a series of presses in one fluid motion. What makes this transformation possible? How do we learn complex dexterous skills such as typing or playing the piano?

Since the early 1950s the discrete sequence production (DSP) task (Fig. 1A) has been used as model to study the acquisition of skilled motor sequences (see Abrahamse et al., 2013 for a recent review). Participants are typically presented with visual stimuli (e.g., numbers or other symbols) and asked to execute the corresponding responses (e.g., finger presses or reaches) as quickly and accurately as possible. While it is known that systematic behavioral training improves performance in sequence execution, the exact processes involved in these behavioral changes remain elusive (Diedrichsen and Kornysheva 2015; Doyon et al. 2018; Wong et al. 2015b).

**Figure 1.**
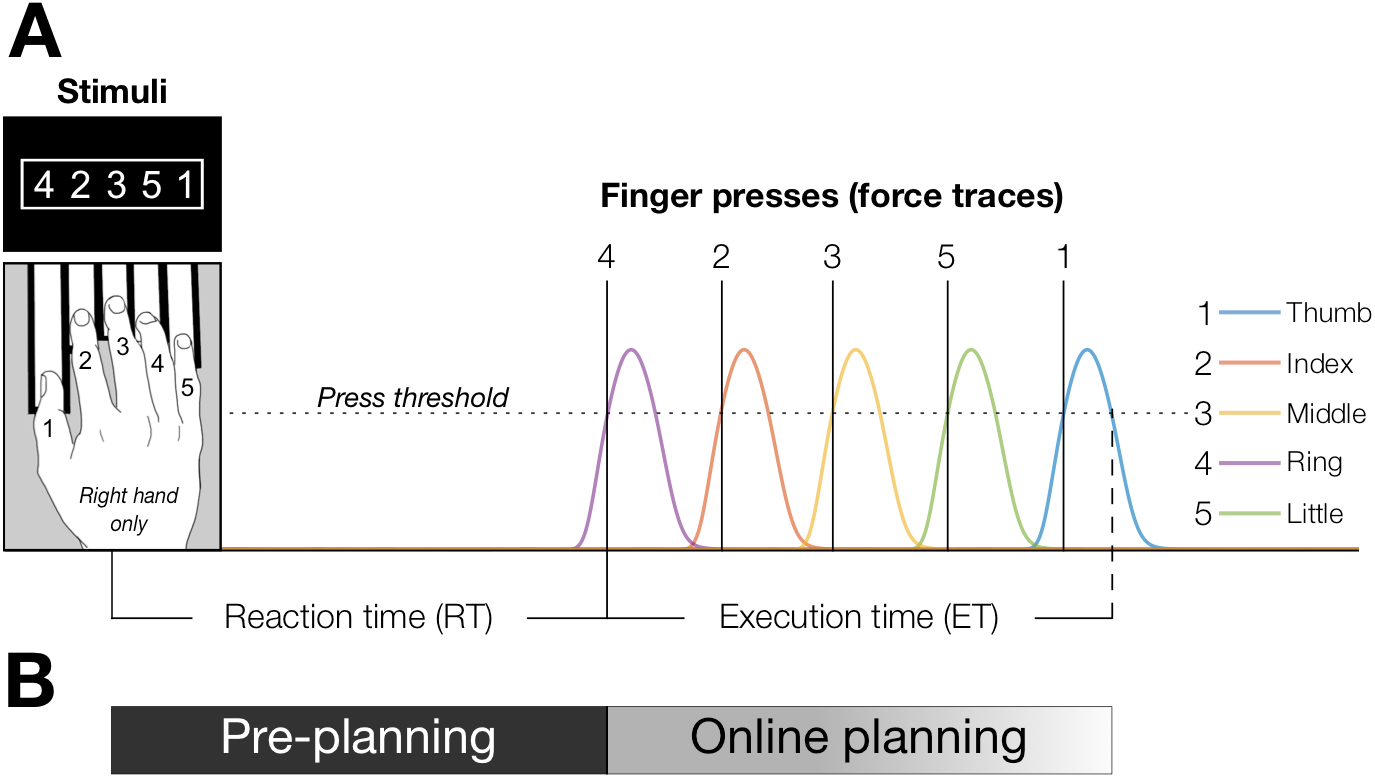
Discrete sequence production (DSP) task. **A.** Visual (e.g., numbers) stimuli instruct which finger presses have to be executed in what order (left-to-right). After a reaction time, five force presses are executed. **B.** *Pre-planning* takes place during the reaction time (i.e., before movement onset); *online planning* occurs in parallel with motor sequence execution (i.e., after movement onset).

Here we sought to characterize the components that underlie the performance improvements in sequence production. Specifically, we attempted to separate processes that occur prior to the start if the sequence, and processes that occur during the execution of the sequence (Fig. 1B).

The first component that we considered is that participants may become better at identifying a stimulus (S; e.g., a number) and selecting the correct response (R; e.g., a finger press). With training, participants could become quicker and more accurate at performing this mapping (Haith et al. 2016; Hardwick et al. 2017), which would speed up both their reaction time and sequence execution. We will collectively refer to improvements in stimulus identification or stimulus-response (S-R) mapping as *single-item selection*.

While it is possible to perform sequences by selecting and executing one action at a time, participants may also pre-select multiple elements prior to execution. This *pre-planning* of multiple elements would make sequence production faster by removing the requirement of action selection during the execution phase. Note that for the purpose of this paper, we do not clear clearly distinguish between selection and planning per se. While planning often implies the specification of movement details beyond merely identifying it, our experimental paradigm does not allow us to make this distinction. Instead, here we will use *selection* to refer to the selection and planning of one element at a time, and *pre-planning* as the process of selecting and planning multiple movements in parallel.

If pre-planning of the full sequence is not possible (e.g., due to the sequence being too long, or preparation time being too short), participants must be able to plan the rest of the sequence ‘on the go’. Thus action selection processes would need to run in parallel to the execution of the first elements. We will refer to this component as *online planning*.

Finally, motor sequence learning can take place at the level of execution processes that are unrelated to selection or planning (Diedrichsen and Kornysheva 2015; Shmuelof et al. 2012; Wong et al. 2015b). This could manifest as an improved ability to press and quickly release a single key, or to transition from one pre-planned key press to the next. We call this component *motor execution*.

To dissociate the influence of these different components to overall speed improvements in motor sequence production, we trained participants using 10 5-item sequences of finger presses. We combined the DSP task with a forced-response paradigm (Ghez et al., 1997; Haith et al., 2016, see Methods section) that enabled us to experimentally manipulate the amount of time to pre-plan a given sequence. We thus investigated how many items participants could pre-plan in advance, and how long this process took. We also established the speed of single-item selection using the same forced-response paradigm in a single-response task. Overall our experiment allowed to break down motor sequence performance into its component processes, and to investigate how each process changed with learning.

## Methods

### Participants

Twenty right-handed volunteers (5 male, 15 female; age range 18-33, average 22.30 years, SD = 4.24) participated in the *Training experiment* (see Experiment paradigm section below) in exchange of monetary compensation. Handedness was verified using the Edinburgh Handedness Inventory (average score 87.25, SD = 14.46). Fifteen of them (4 male, 11 female; average age 22.47 years, SD = 4.67) participated in the *Retention test* (see Experiment paradigm section below) roughly three months after the last training day (average time difference 82 days, SD = 15.16). The experimental procedures were approved by the local ethics committee at Western University (London, Ontario, Canada), and all participants provided written informed consent. None of the participants was a professional musician (average musical experience 3.88 years, SD = 3.82), nor had any history of neurological disorder.

### Apparatus and stimuli

Sequences of finger presses were executed on a piano-like keyboard device (as shown in Fig. 1A) comprised of 5 keys equipped with force transducers (FSG-15N1A, Sensing and Control, Honeywell; dynamic range, 0-25 N) that measured isometric force presses exerted by each finger with an update rate of 2 ms. To account for sensor drifts, a zero-force baseline was recalibrated at the beginning of each block. A key press was recognised when the sensor force exceeded a press threshold of 1 N (e.g., vertical solid lines in Fig. 1A). A key was considered released when the force returned below 1 N (e.g., vertical dashed line in Fig. 1A). The device was custom built and has been described in detail previously (Kornysheva and Diedrichsen 2014; Wiestler and Diedrichsen 2013; Yokoi et al. 2017, 2018). The stimuli consisted of white numeric characters on black background framed by a white rectangle (as shown in Fig. 1A) presented on a computer display (stimuli height 1.5 cm; visual angle ~2°).

### DSP Task

A discrete sequence production (DSP) task required participants to make isometric force presses with the fingers of their right hand. The instructing stimulus was a five-item sequence cue (a string of numbers ranging 1 to 5) which indicated, from left to right, the order in which the fingers had to be pressed (e.g., 1 = thumb, 5 = little finger). Once present on the screen, the stimuli remained visible for the entire duration of the trial, thus participants had always full explicit knowledge of the sequence of finger presses to be executed. Participants were instructed to complete the sequence as quickly and accurately as possible. Performance was evaluated in terms of both speed and accuracy in sequence production. Speed was quantified in execution time (ET), defined as the time from the first key press (Fig. 2A, P1) to the release of the last key press (Fig. 2A, black dashed vertical line). With each press, the corresponding white number turned either green for correct responses, or red for erroneous responses (Fig. 2A, top). A single press error in the sequence, or timing error on the first press, invalidated the whole trial, so accuracy was calculated as percent error rate (ER) per block of trials (i.e., number of error trials / number of total trials × 100). In case of an error, participants were instructed to complete the sequence and move on to the next trial. After completing the entire sequence (Fig. 2A, black dashed vertical line), performance points appeared on the screen, replacing the sequence cues (Fig. 2A, top right; see Feedback section below) during 500 ms of inter-trial interval (ITI) before moving on to the next trial.

**Figure 2.**
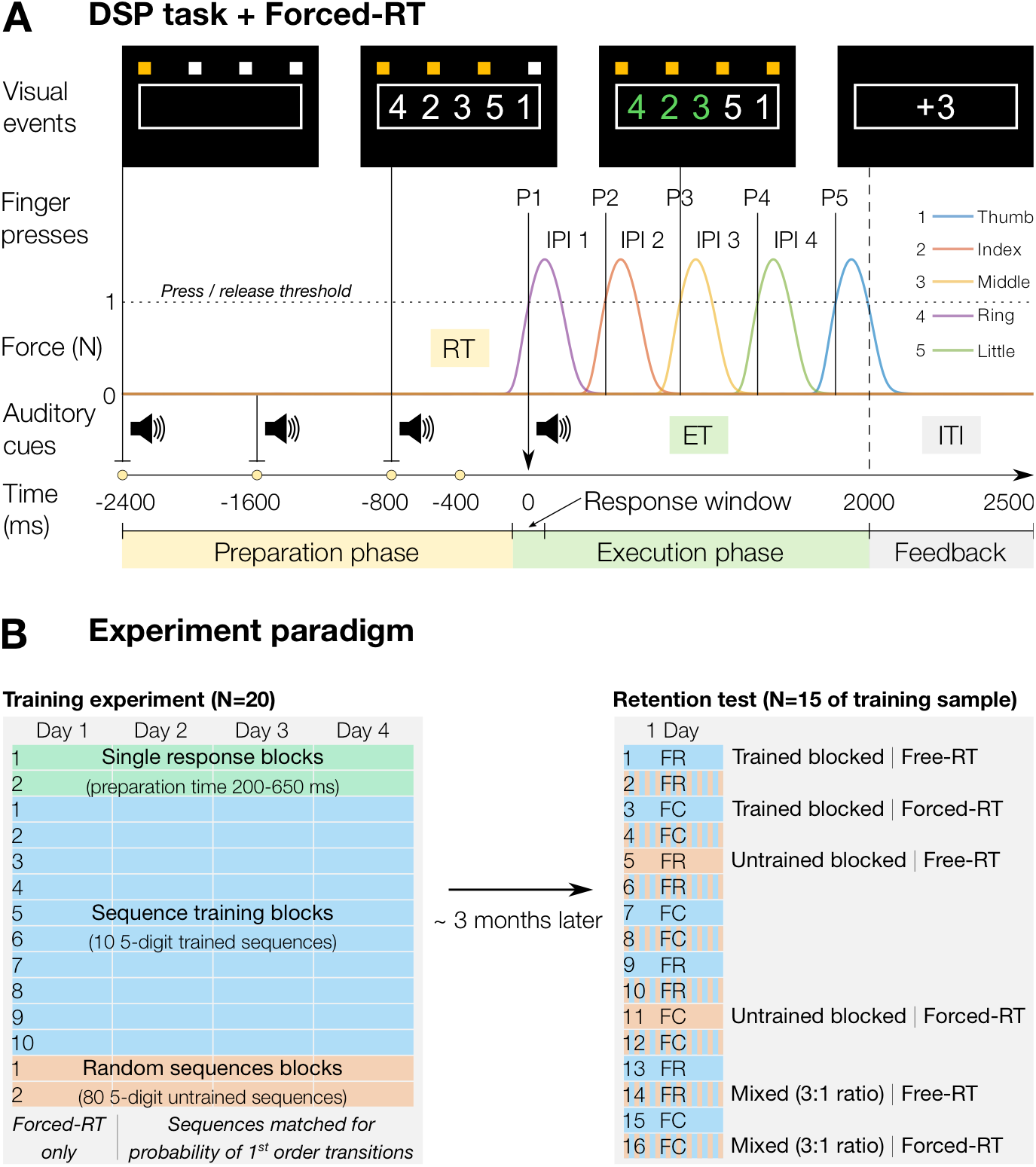
Experimental task and paradigm. **A.** A series of 4 audio-visual signals (800 ms apart) specifies the timing of movement initiation (vertical arrow) within the acceptable response window. Sequence cues appear at one of four time points (yellow dots) before the 4^th^ signal (Preparation phase). After completing the sequence (Execution phase), participants receive points depending on their performance. Colored lines illustrate schematic force traces for the 5 finger presses in a sequence. The horizontal dotted line denotes the force threshold for a press/release. Vertical lines indicate press onsets (P1 = first press; IPI 1 = first inter-press interval). The vertical dashed line represents the release of the last press (end of the Execution phase). RT = reaction time, ET = execution time, ITI = inter-trial interval. **B.** *Left*: the Training experiment (N = 20) consisted of 4 days of training on single response (green), sequence training (blue), and random sequences (orange) blocks. *Right:* after ~3 months we called the same participants back for a Retention test (N = 15) with blocks of trials including either trained (blue), untrained (orange), or mixed trained/untrained (striped) sequences. Mixed blocks contained 30 trained and 10 untrained sequences presented in random order. We also alternated the Forced-RT (FC) and Free-RT (FR) paradigm every two blocks. The initial paradigm was counterbalanced across participants.

### Forced-RT paradigm

In order to manipulate sequence preparation time we combined the DSP task with a forced reaction time (Forced-RT) paradigm (e.g., Ghez et al., 1997; Haith et al., 2016).

On each trial, a fully predictable series of four regularly-paced auditory tones (800 ms apart; Fig. 2A, bottom) cued the time of movement initiation (Fig. 2A, black vertical arrow). Participants were trained to press the first finger of the sequence as synchronously as possible with the 4^th^ tone. The informative sequence cue (i.e., the numbers on the screen) could appear 400, 800, 1600, or 2400 ms before the 4^th^ tone (yellow dots in Fig. 2A), giving participants variable time to plan the whole sequence before making the first press. To increase the salience of the Forced-RT paradigm, the tones were accompanied by visual cues (four horizontally-aligned white squares on black background, each turning yellow in synchrony with an auditory tone; Fig. 2A, top). A response window of 200 ms centered on the 4^th^ tone (i.e., up to 100 ms before and 100 ms after the tone) defined the acceptable range of response times for the first press to count as correct timing. In case of a timing error, participants received immediate visual feedback, with the words *“Early”*, or “*Late”*, appearing on the screen above the sequence cue. This paradigm enabled us to manipulate the amount of time available for sequence pre-planning (Fig. 1B), and indirectly assess the amount of sequence planning by forcing participants to start the sequence at faster-than-normal RTs, while measuring their performance as a function of the imposed preparation time (Haith et al. 2016).

For single response blocks (Fig. 2B; see Experiment Paradigm section below), participants were presented with five numbers but asked to only focus on the first (a number from 1 to 5), ignore the other four (filler numbers from 6 to 9 and 0), and produce the corresponding single finger press in synchrony with the 4^th^ tone. Because single-item selection is faster than sequence pre-planning, the preparation times in the single response blocks ranged from 200ms to 650ms before the 4^th^ tone in 10 steps of 50 ms.

### Feedback

In order to motivate participants to improve further in sequence production speed once they became comfortable with the task, we provided them with feedback about their performance throughout each training and testing session.

At the end of each trial, participants received a performance score according to the following point system: −3 points for large timing errors (i.e., more than 300 ms before, or after, the last tone); −1 points for small timing errors (i.e., between 300 and 100 ms before, or after, the last tone); 0 points for correct timing (i.e., less than 100 ms before, or after, the last tone) but wrong finger press; +1 points for correct timing and press, but ET 20% or more higher than upper ET threshold; +2 points for correct timing and press, but ET between upper and lower ET thresholds; and +3 points for correct timing and press, and ET 5% or more lower than lower ET threshold. Upper and lower ET thresholds determining the performance score would decrease from one block to the next if both of the following performance criteria were met: median ET in the last block faster than best median ET so far in the session, and mean ER in the last block < 30%. If either one of these criteria was not met, the thresholds for the next block remained unchanged (i.e., the same as previous block). At the end of each block of trials, the median ET, mean ER, and points earned were displayed to the participants.

The point system remained identical for Single response blocks (Fig. 2B; see Experiment Paradigm section below), with the exception that there was no sequence execution speed (i.e., no ET), but only finger selection accuracy. Thus making the correct finger press with correct timing directly produced +3 points. Given the shorter preparation time allowed in Single response blocks, we warned participants that the task was supposed to be challenging and that, if they felt like they didn’t have enough time to plan the correct press, they should randomly choose which finger to press, as it would be more beneficial for them to guess wrong at the right time (0 points), than to respond correctly with incorrect timing (negative points).

### Experiment paradigm

The present study was composed of 2 parts: a *Training experiment* and a *Retention test* (Fig. 2B).

#### Training experiment

The main purpose of the training experiment was to examine the effects of training and preparation time (and their interaction) on single response selection and sequence production. Sequence training was manipulated by using the DSP task and two non-overlapping sets of 5-item sequences: a set of 10 repeating sequences (trained sequences; each sequence presented 40 times per day, in random order), and a set of 80 novel sequences (untrained sequences; each sequence only presented once per day, in random order). The two sets of sequences were matched for probability of 1^st^ order finger transitions, i.e. how often a specific finger followed another specific finger. Thus, differences between trained and untrained sequences cannot be attributed to simple differences in learning such 1^st^ order transitions. Each testing session of the training experiment (one per day over 4 days) consisted of 14 blocks of trials performed in the following order: 2 Single response blocks (50 trials each), 10 Sequence training blocks (40 trials each, trained sequences only), and 2 Random sequences blocks (40 trials each, untrained sequences only). Single response blocks were intended to assess eventual performance improvements in single finger selection accuracy as a function of preparation time. Sequence training blocks served to train participants on the finger movements associated with each sequence in the set of trained sequences (i.e., to develop sequence-specific learning). Random sequences blocks were used to test sequence production performance on a novel set of untrained sequences.

#### Retention test

Roughly three months after the last session of the training experiment, we called participants back for a retention test. The main purpose of this retention test was to check 1) whether the components of sequence learning identified in the training experiment were stable or susceptible to forgetting, and 2) whether the results obtained in the training experiment would replicate even when using a Free-RT paradigm (i.e., no time pressure requirements to start the sequence), or when mixing trained and untrained sequences within the same block. The retention test consisted of 16 blocks of sequence production in which we alternated the Forced-RT with a Free-RT paradigm every two blocks (the first paradigm type, whether Free- or Forced-RT, was counterbalanced between subjects). Each block could be either a block of trained sequences (3 blocks per paradigm, 40 trials each), a block of untrained sequences (1 block per paradigm, 40 trials each), or a mixed block (4 blocks per paradigm, 40 trials each: 30 trials of trained sequences, and 10 trials of untrained sequences randomly interleaved), following the order shown in Fig. 2B (right panel). The sets of trained and untrained sequences, the preparation time levels, and the point system for feedback on performance remained identical as in the training experiment. Additionally, to test for explicit sequence memory retention, both before and after the sequence production task, we administered a questionnaire that included both free sequence recall (e.g., *“fill in the numbers for each of the 10 repeating sequences – if you don’t remember, guess’)*, and sequence recognition tests (e.g., *“for each of the following 30 sequences, indicate whether it was one of the repeating, or novel, sequences – if unsure, guess’)*.

### Data analysis

All data were analyzed offline using custom code written in MATLAB (The MathWorks, Inc., Natick, MA). To assess behavioral improvements in the Single response task, for each subject, and training day, we calculated the mean finger selection accuracy as a function of preparation time. For this analysis we considered only trials in which the timing had been correct (i.e., press within the response window). To quantify improvements in mean finger selection accuracy (an indirect measure of single-item selection speed), we fitted the data from each training day to the following logistic function, separately for each subject:

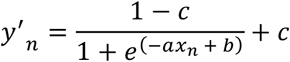

where *y*′*_n_* is the predicted finger selection accuracy for trial n; *x_n_* is the manipulated preparation time; *a* and *b* are free parameters determining, respectively, the steepness of the function at the midpoint, and the location of the midpoint on the x-axis (i.e., the preparation time where the midpoint occurs). The offset *c* = 0.2 constrained the logistic function to range between 0.2 (chance selection accuracy) and 1. The parameters *a, b* were then fitted to the data of each day separately using MATLAB’s *fminsearch* routine to minimize the Bernoulli loss function:

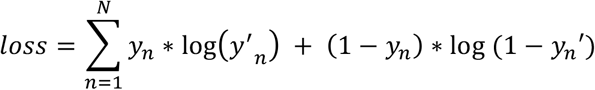

where *y* is the observed and *y*′ the predicted selection accuracy. Due to poor performance in single response blocks and insufficient number of trials for some of the conditions, the data from one subject could not be fitted and was therefore excluded from this analysis (thus, N = 19).

Statistical analyses to assess sequence-general and sequence-specific learning, and improvements in sequence pre-planning speed included two-tailed paired samples t-tests and a within-subject repeated measures ANOVA with factors training and preparation time. Error trials (both timing and press errors) were excluded from this analysis. Additionally, as a visual aid to the interaction between preparation time and training, we plotted the ET of trained and untrained sequences in terms of percentage of the ET for the longest preparation time (2400 ms). This allowed us to show the benefit of longer preparation times as gain in sequence execution speed while directly comparing trained and untrained sequences.

Finally, for each participant, explicit sequence-memory retention was quantified as mean free sequence-recall accuracy, and mean *d-prime* (*d*’) was taken as a measure of sensitivity in sequence recognition.

## Results

### Training leads to sequence-specific and sequence-general learning

Improvements in speed were observed both for trained, as well as for novel, untrained sequences. From the first to the last training day, participants improved their average execution time from 1564 ± 131 ms to 1288 ± 94 ms even on untrained sequences (*t*_19_ = 5.03, *p* < 0.0001; Fig. 3A), demonstrating sequence-general learning. To determine the amount of sequence-specific learning, we compared the performance on the 2 untrained blocks in the end of each day to the 2 preceding blocks of trained sequences. On the fourth day, we observed that participants executed trained sequences significantly faster than untrained sequences (trained: 958 ± 75 ms, untrained: 1288 ± 94 ms, *t*_19_ = 9.64, *p* < 0.0001; Fig. 3A), while maintaining comparable accuracy levels (trained: 77.5 ± 2.5 %; untrained: 79.3 ± 2.2 %, t19 = 0.96, *p* = 0.35; Fig. 3B).

**Figure 3.**
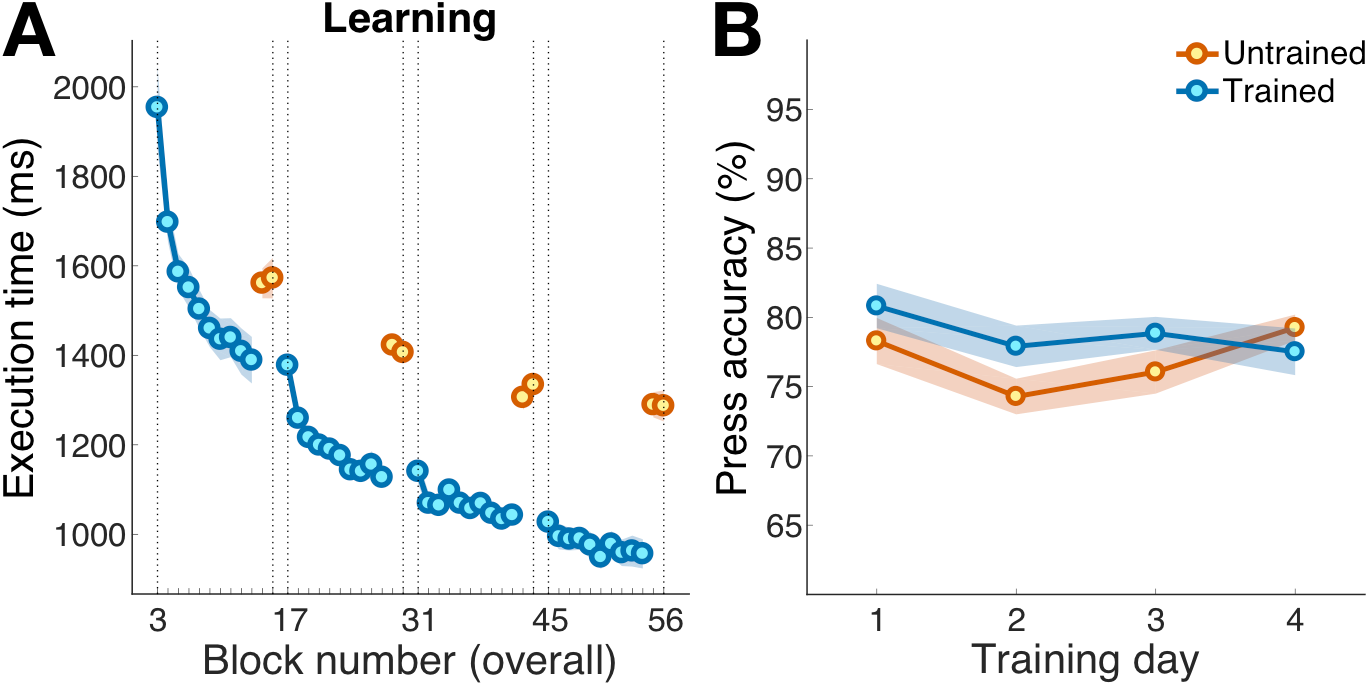
Sequence-general and sequence-specific learning. **A.** Mean sequence execution time plotted as a function of block number, separately for trained (blue) and untrained (orange) sequences. Vertical dotted lines demarcate the beginning and the end of each testing session (one per day). Shaded areas indicate within-subject standard error. **B.** Mean sequence execution accuracy (among trials with correct timing) as a function of training day, separately for trained (blue) and untrained (orange) sequences. Shaded areas indicate within-subject standard error.

Thus, in addition to the substantial improvement even for the execution of untrained sequences, participants showed a 330 ms advantage for trained sequences.

### Single-item selection becomes quicker with practice

One factor that may underlie sequence-general learning effects is an improved ability to associate each number with the corresponding finger. We measured this ability using a forced-response paradigm on single-item selection. Consistent with previous results (Haith et al. 2016; Hardwick et al. 2017), we found that participants were able to select the correct finger with increasing probability for shorter preparation times (Fig. 4A). Fitting a logistic function to these data allowed us to quantify learning in terms of both selection accuracy and speed (Fig. 4B). Given a preparation time of 400ms, single finger selection improved by 14.48 % from day 1 to day 4 of testing (*t*_18_ = −3.72, *p* = 0.0016). Correspondingly, the average preparation time required to reach at least 80% finger selection accuracy decreased by 100 ± 26 ms (*t*_19_ = 3.88, *p* = 0.0011).

**Figure 4.**
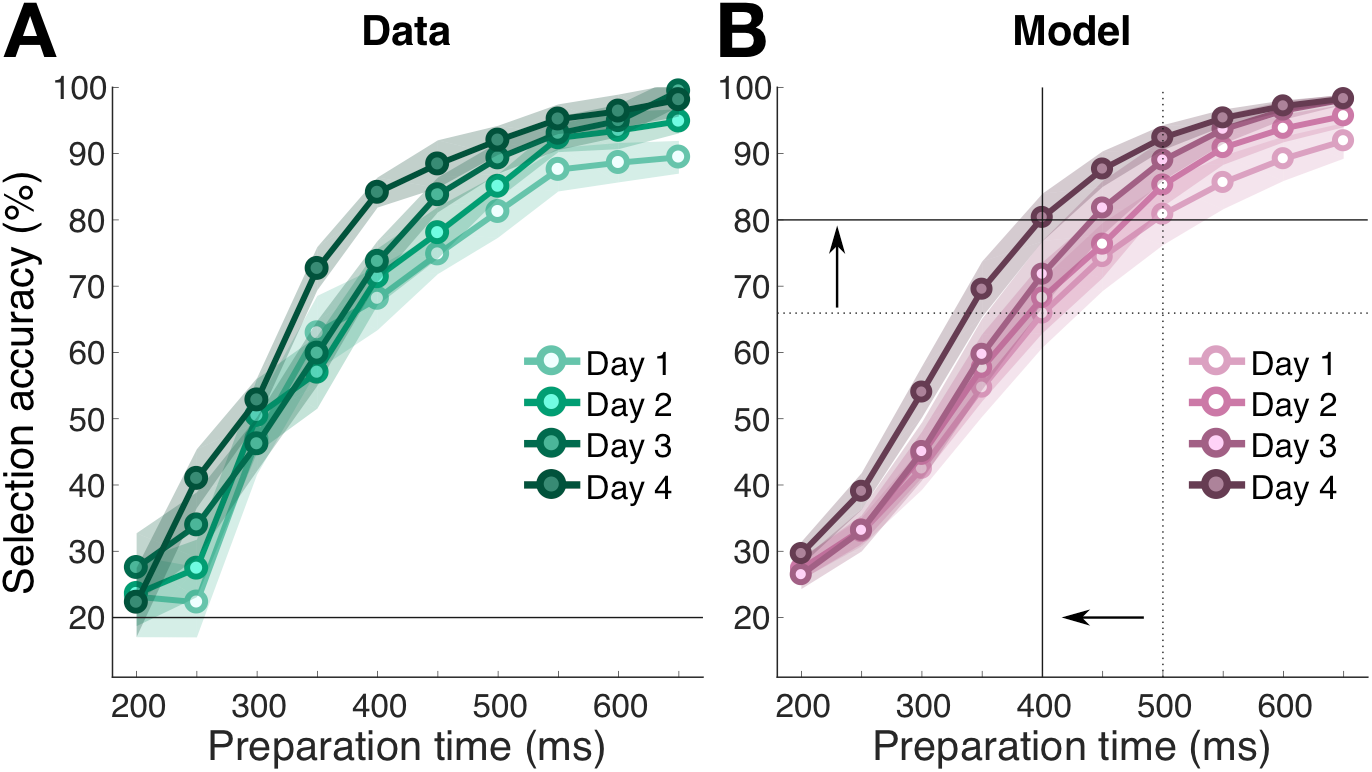
Selection of individual action elements improves with practice. **A.** Mean single finger selection accuracy plotted as a function of preparation time (determined by Forced-RT), separately for each testing day. The horizontal solid line denotes chance level for selection accuracy. Shaded areas indicate standard error of the mean. **B.** Logistic function model fits for data shown in A, separately for each testing day. Solid lines and arrows serve as a visual aid to appreciate performance improvements in selection accuracy given a 400 ms preparation time (vertical solid line), and improvements in selection speed in order to reach 80 % selection accuracy (horizontal solid line) from day 1 (dotted lines) to day 4. Shaded areas indicate standard error of the mean.

These results indicate that learning effects in sequence production can be partly explained by improvements in the selection and execution of individual sequence elements. Indeed, a ~100 ms speed improvement per digit could potentially fully account for the observed improvements for untrained sequences. However, it cannot account for the added performance benefit for trained sequences, as the speed-up in single digit selection would benefit both trained and untrained sequences equally.

### Participants pre-plan the first 3 items in a sequence

Although sequence pre-planning cannot be measured directly at the behavioral level (Haith et al. 2016), having more time to plan a sequence should result in better (i.e., faster, more accurate) sequence execution. Therefore, we used a forced-response paradigm to infer how much of a sequence had been pre-planned. Indeed, we found that longer preparation times led to faster sequence execution (Fig. 5A), in any phase of training. This was confirmed by a within-subject ANOVA that included the last 2 trained and the last 2 untrained blocks of each day: averaged across day and sequence type (i.e., trained or untrained), the main effect of preparation time on ET was significant (*F*_3,57_ = 58.63, *p* < 0.0001). In other words, with more time to prepare sequences are planned better and executed faster.

**Figure 5.**
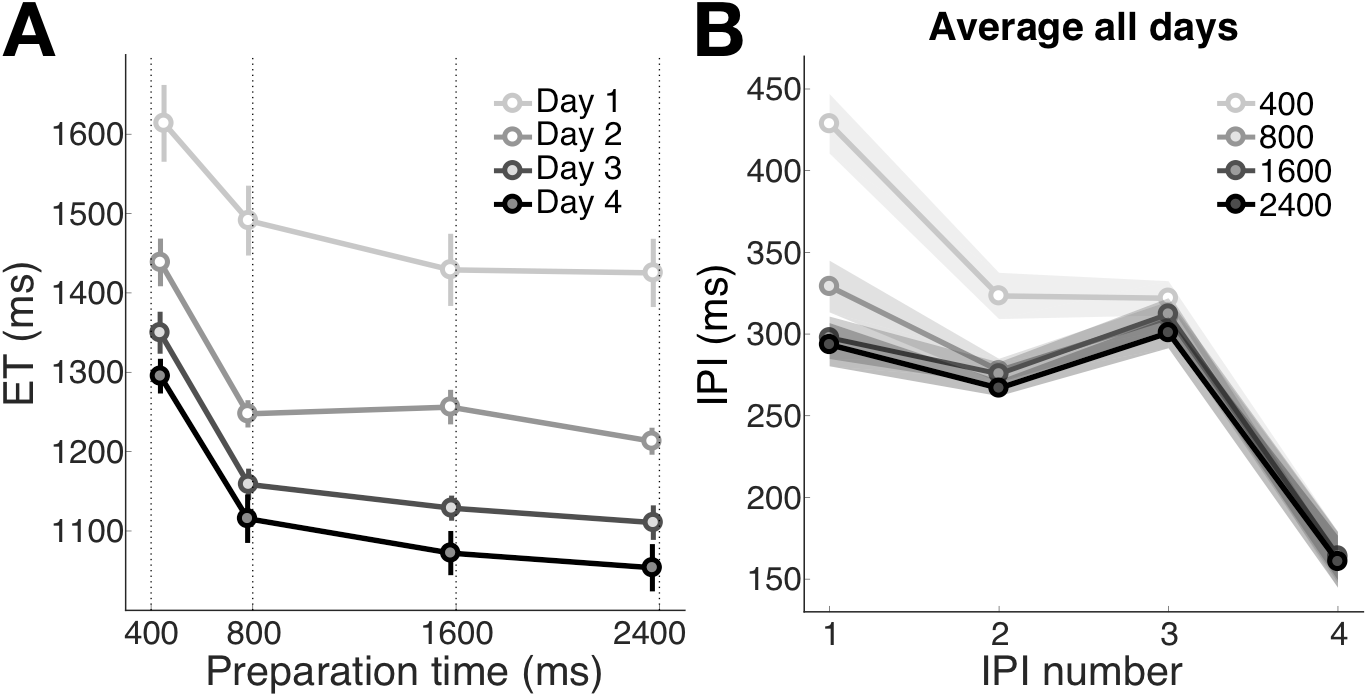
Preparation time affects the quality of sequence pre-planning. **A.** Mean ET as a function of preparation time. Separate lines indicate results for each training day (different shades of gray). Results are averaged across trained and untrained sequences, using the last 4 blocks for each training day. Error bars reflect standard error of the mean. For each preparation time, the actual average RT is shown, with the instructed RT indicated by the dotted line. **B.** Mean duration for each inter-press interval (IPI), averaged across training days, with separate lines for each preparation time level. As in A, results are averaged over the last 4 blocks of each day, collapsing across trained and untrained sequences. Shaded areas indicate within-subject standard error of the mean.

The results of the single response task indicate that 400 ms are enough to select and plan the first item in the sequence. Therefore, 2400 ms should be enough time to fully select and plan a 5-item sequence. But how many steps ahead do participants actually pre-plan – i.e., what is their pre-planning *capacity*? To answer this question, we inspected the profile of inter-press intervals (IPIs), averaged across training days and trained or untrained sequences (last 4 blocks from each day), separately for each preparation time level (Fig. 5B). We observed that the 1^st^ IPI took 100 ± 14 ms longer in the 400 ms than in the 800 ms preparation time condition, indicating that participants were making use of the extra preparation time to pre-plan the transitions between the 1^st^ and the 2^nd^ finger press (*t*_19_ = 6.91, *p* < 0.0001). The difference between 400 ms and 800 ms preparation time condition was also significant for the 2^nd^ IPI (mean difference = 46 ± 10 ms; *t*_19_ = 4.41, *p* = 0.0003) and 3^rd^ IPI (mean difference = 12 ± 5 ms; *t*_19_ = 2.46, *p* = 0.02). In contrast, no effect of preparation time on IPI duration was found for the 4^th^ IPI (*F*_3,57_ = 0.23, *p* = 0.87). This suggests that, even for the longest preparation times, participants did not pre-plan the entire sequence. Longer preparation times only resulted in faster execution of the first three elements in the sequence (an indirect estimate of sequence pre-planning), but did not affect the execution speed of the last two elements (i.e., these elements were not pre-planned in advance).

Taken together, our results provide evidence that on average participants preplan at least 3 elements into the future. It follows that, even a short 5-element sequences could not be fully pre-planned, regardless of the given amount of preparation time. As the remaining elements still need to be selected and planned, this type of planning has to occur online, i.e., while already executing the beginning of the sequence.

### Sequence-specific learning speeds up pre-planning, does not increase preplanning capacity

Next we asked how much the ability to pre-plan improves with training. To investigate the influence of sequence-specific learning on pre-planning speed, we compared the effects of varying preparation time on ET for trained and untrained sequences at day 4 (Fig. 6A, solid lines). While we observed the general decrease in ET with longer preparation times (*F*_3,57_ = 40.32, *p* < 0.0001), trained sequences were always faster than untrained sequences (*F*_1,19_ = 107.62, *p* < 0.0001), even for the longest preparation time (mean difference = 283 ± 38 ms; *t*_19_ = 7.35, *p* < 0.0001). This suggests that even if pre-planning has reached full capacity, there is a substantial advantage for extensively trained sequences. This advantage might reflect learning to execute coordinated transitions between finger presses or improvements in online planning (see below).

**Figure 6.**
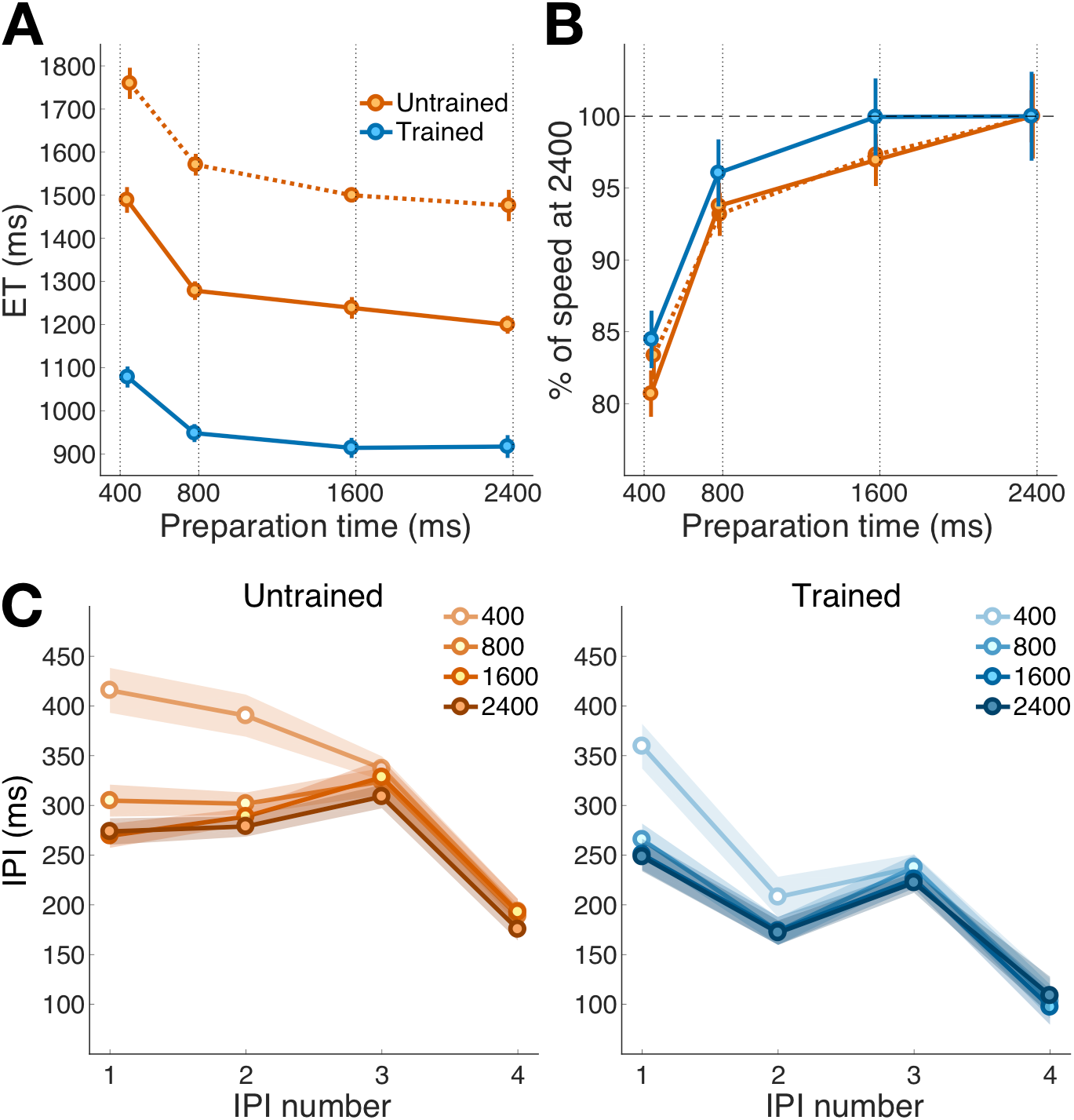
Training makes pre-planning faster, but not more complete. To assess sequence-general learning we compared performance for untrained sequences between day 1 and day 4. To assess sequence-specific learning, we compared trained and untrained sequences on day 4 (the last 4 blocks: 2 trained, 2 untrained). **A.** Mean ET as a function of preparation time, separately for untrained sequences on day 1 (orange dotted), trained (blue) and untrained (orange solid) sequences on day 4. Other figure conventions are the same as in Figure 5A. **B.** Speed (1/ET), expressed as percentage of the individual speed reached at 2400 ms preparation time, separately for untrained sequences on day 1 (orange dotted), trained (blue) and untrained (orange solid) sequences on day 4. **C.** Mean interpress interval (IPI) duration on day 4, separately for untrained (left) and trained (right) sequences, and preparation time levels. Shaded areas indicate standard error of the mean.

Importantly, however, longer preparation times benefitted the execution of untrained sequences more than the trained ones, as confirmed by a significant interaction between preparation time and sequence type (*F*_3,57_ = 4.17, *p* = 0.009). To visualize this effect better, we normalized the speed for each condition (1/ET) by the speed reached given the longest preparation time (2400 ms; Fig. 6B, solid lines). This analysis shows that participants could reach pre-planning capacity for trained sequences in 1600 ms, whereas they still benefitted from additional time to pre-plan the untrained sequences. This result indicates that sequence-specific learning makes the pre-planning of known action sequences faster.

In contrast, sequence-general learning did not seem to improve the speed of pre-planning. We compared the performance curve for untrained sequences between day 1 (Fig. 6A, dotted lines) and day 4. Despite the clear improvement in sequence execution speed from day 1 to day 4 for untrained sequences (*F*_1,19_ = 10.40, *p* = 0.004), we found no significant interaction between preparation time and day (*F*_3,57_ = 0.19, *p* = 0.91). After normalization to the fully-prepared execution (2400ms preparation time), the speed of preplanning remained stable (Fig 6B, dotted lines). Together these analyses reveal that improvements in pre-planning speed are present for trained, but not for untrained sequences.

We then inspected the IPI profiles of trained and untrained sequences on day 4 (Fig. 6C) to ask whether sequence-specific learning would also increase preplanning capacity. This analysis confirmed that on average participants can pre-plan at least 3 elements into the future (effect of preparation time on 2^nd^ IPI, even for trained sequences: 400 ms condition vs. 800 ms condition, t(19) = 2.14, p = 0.04). However, the lack of an effect of preparation time on the 4^th^ IPI of trained sequences (400 ms condition vs. 2400 ms condition; *t*_19_ = −0.17, *p* = 0.87) suggests that, even provided the longest preparation time, pre-planning was not complete. This means that, despite the improvements in pre-planning speed that resulted from extensive training, pre-planning capacity did not substantially improve.

At the same time, even the last IPI was executed substantially faster for trained as compared to untrained sequences (mean difference = 67 ± 11 ms; *t*_19_ = 5.94, *p* < 0.0001). From this we must conclude that improvements in online planning of the remaining elements play a significant role in the behavioral improvements observed in sequence production.

### Retention test: learning effects are robust to experimental variations

The results of the training experiment suggest that improvements in single-item selection, pre-planning, and online planning drive the development of skilled sequence performance. To show that the observed effects are stable across time, and that they are not dependent on how sequence performance was tested, we conducted a retention test about 3 months after the training experiment.

First, we sought to test whether the sequence-specific advantages could arise from a general motivational effect given by the fact that in the training experiment trained and untrained sequences were executed in blocked fashion. Knowledge of being in a Sequence training block might have acted as a reward, increasing participants’ motivation, thus leading to performance improvements (Wong et al. 2015b). Therefore, we tested trained and untrained sequences under mixed and blocked conditions (Fig. 7). We replicated the main effects of preparation time (*F*_3,42_ = 34.12, *p* < 0.0001), sequence type (*F*_1,14_ = 89.28, *p* < 0.0001), and the significant interaction between preparation time and sequence type on mean ET (*F*_3,42_ = 3.38, *p* = 0.027). The fact that trained sequences were executed faster than untrained sequences in the retention test suggests that participants had some memory of the trained sequences. Notably, for both blocked (Fig. 7A) and for mixed (Fig. 7C) presentation of trained and untrained sequences, we could confirm that there was a benefit of sequence-specific training even for fully prepared sequence (2400ms preparation time, blocked: 131 ± 36 ms, *t*_14_ = 3.64, *p* = 0.0027; mixed: 131 ± 29 ms, *t*_14_ = 4.55, *p* = 0.0005). Hence the superior performance on trained compared to untrained sequences cannot be explained by a difference in overall motivation between blocks.

**Figure 7.**
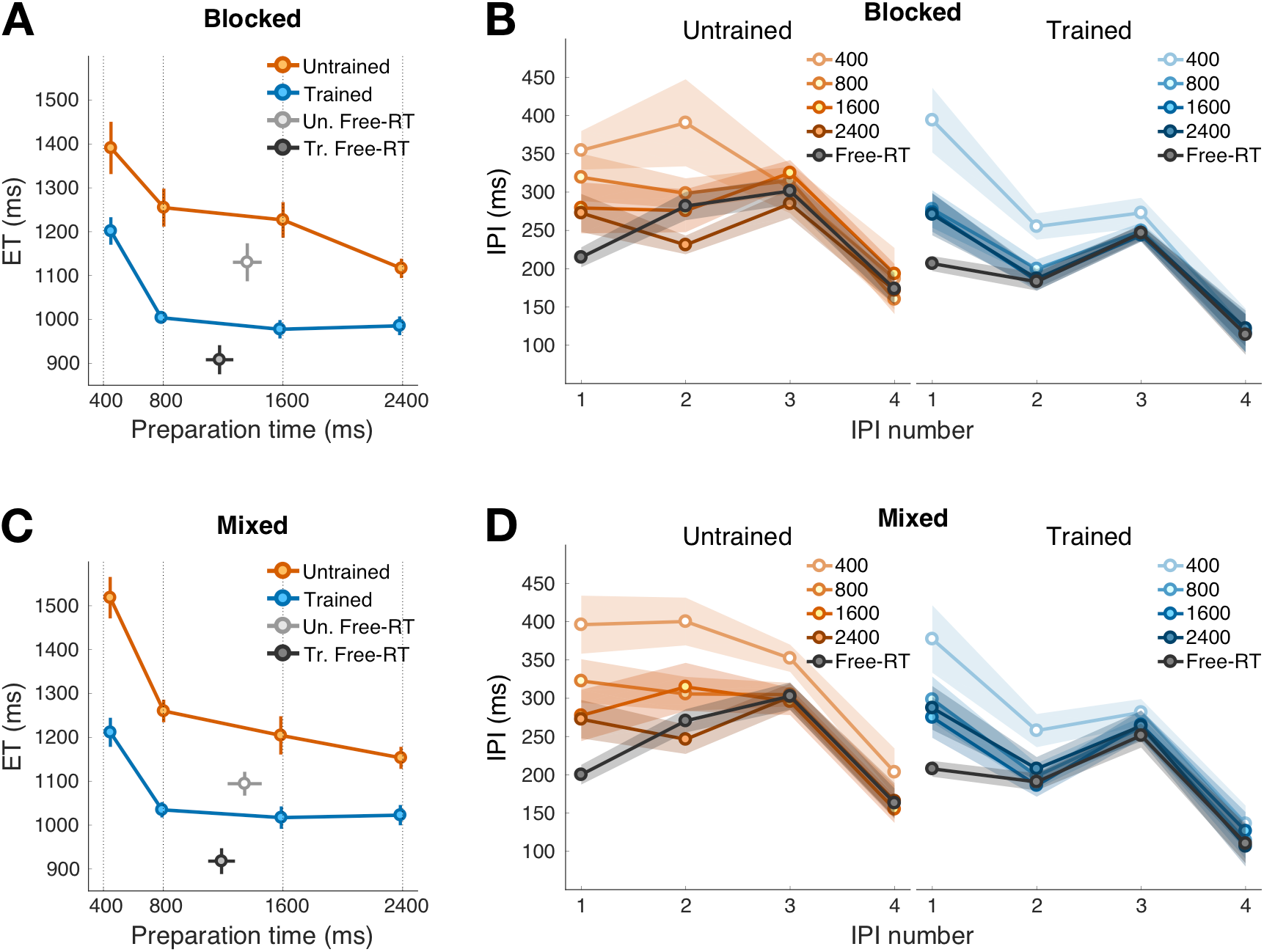
Performance improvements are stable across time and experiment paradigm (Retention test). **A.** Mean ET as a function of preparation time levels with the factor sequence type Blocked, separately for trained (blue), untrained (orange) sequences in the Forced-RT condition, and trained (dark gray), untrained (light gray) sequences in the Free-RT condition. Other figure conventions are the same as in Figure 5A. **B.** Mean duration for each inter-press interval (IPI) in Blocked blocks, separately for untrained (left) and trained (right) sequences, and preparation time (different shades of, respectively, orange and blue). For sake of comparison, mean IPI durations for the Free-RT task are plotted in gray. Shaded areas indicate within-subject standard error of the mean. **C.** Same as A, but with factor sequence type Mixed. **D.** Same as B, but for Mixed blocks of trials.

Second, the retention test allowed us to test whether the results obtained in the training experiment with a Forced-RT paradigm would replicate when participants are free to start the sequence at will. The analysis of the Free-RT blocks (gray data points in Fig. 7A-7C) confirmed that trained sequences were executed faster than untrained ones (mean ET difference, blocked: 222 ± 45 ms; *t*_14_ = 4.91, *p* = 0.0002; mixed: 177 ± 25 ms; *t*_14_ = 7.01, *p* < 0.0001). Moreover, we found a significant reaction time advantage for trained as compared to untrained sequences in Free-RT blocks (mean RT: 170 ± 54 ms; *t*_14_ = 3.18, *p* = 0.006), suggesting that trained sequences could be pre-planned more quickly. Taken together, these results corroborate the findings of the training experiment, and indicate that these results can be replicated outside of the Forced-RT paradigm.

Interestingly, when allowed to take as much time as needed before starting the sequence, participants had an average RT of ~1300 ms but showed better performance than the 1600 and 2400 ms preparation time conditions in the Forced-RT paradigm: ET in the Free-RT condition was marginally faster than in the Forced-RT condition (mean RT difference across sequence types: 80 ± 37 ms; *t*_14_ = 2.17, *p* = 0.048). This effect likely reflects the dual-task nature of the forced-RT paradigm: participants have to keep track of the audio-visual cues and plan the response at the same time. The additional requirement may hinder sequence planning and execution, thus resulting in slightly poorer performance.

Finally, the retention test allowed us to explore also to what degree sequence-specific learning depends on an explicit memory of the trained sequences. The development of explicit sequence memory might have facilitated conscious retrieval of trained sequences in the retention test, perhaps contributing to the observed performance advantage in sequence planning and execution. However, it should be noted that participants were always provided with full explicit sequence knowledge during the task, as the sequence cues remained visible on the screen throughout the execution of both trained and untrained sequences. So there was no incentive for the participants to memorize the sequences. This became evident from the results of the tests for explicit sequence knowledge retention. Participants performed poorly in both free sequence recall (mean recall out of 10 sequences: presequence production = 12.67 ± 3.16 %; post-sequence production = 23.33 ± 3.33 %; overall = 18.00 ± 2.88 %) and sequence recognition (mead d’: pre-sequence production = 0.65 ± 0.33; post-sequence production = 1.05 ± 0.17; overall = 0.85 ± 0.23). These results indicate that the memory of trained sequences was predominantly implicit.

Overall, the results of the retention test support the idea that the sequence-specific improvements in planning observed in the training experiment did not depend on the particular choice of design, or the presence of dual-task requirements in the forced-RT paradigm.

### Improvements in online planning drive sequence-specific learning

The retention test also allowed us to more closely investigate sequence-specific improvements in online planning. As in the training experiment, we confirmed that participants could still pre-plan an average of 3 elements into the future, and we found no effect of preparation time on the 4^th^ IPI (*F*_3,42_ = 2.65, *p* = 0.06; Fig 7B-7D). Once again, we took this result as a sign that participants could not pre-plan the whole sequence at once, which indirectly implies that online planning must have occurred.

To further investigate the contribution of online planning to sequence-specific learning, we re-analyzed the IPIs from the training and retention experiments, grouping the first two (IPI 1-2) and the last two (IPI 3-4) IPIs in the sequence (Fig. 8). As we have shown, the first two IPIs could be largely pre-planned (as their speed varied with preparation time), whereas the last two IPIs needed to rely on online planning to a much greater extent. Thus, if we found an effect of sequence type on IPI 1-2 at the longest preparation time (i.e., for well pre-planned presses), it would provide evidence that sequence execution processes had improved beyond the advantages of pre-planning. Additionally, if we found a larger effect of sequence type on IPI 3-4 (i.e., for less pre-planned presses), we would take this as evidence that online planning got better following sequence-specific training.

**Figure 8.**
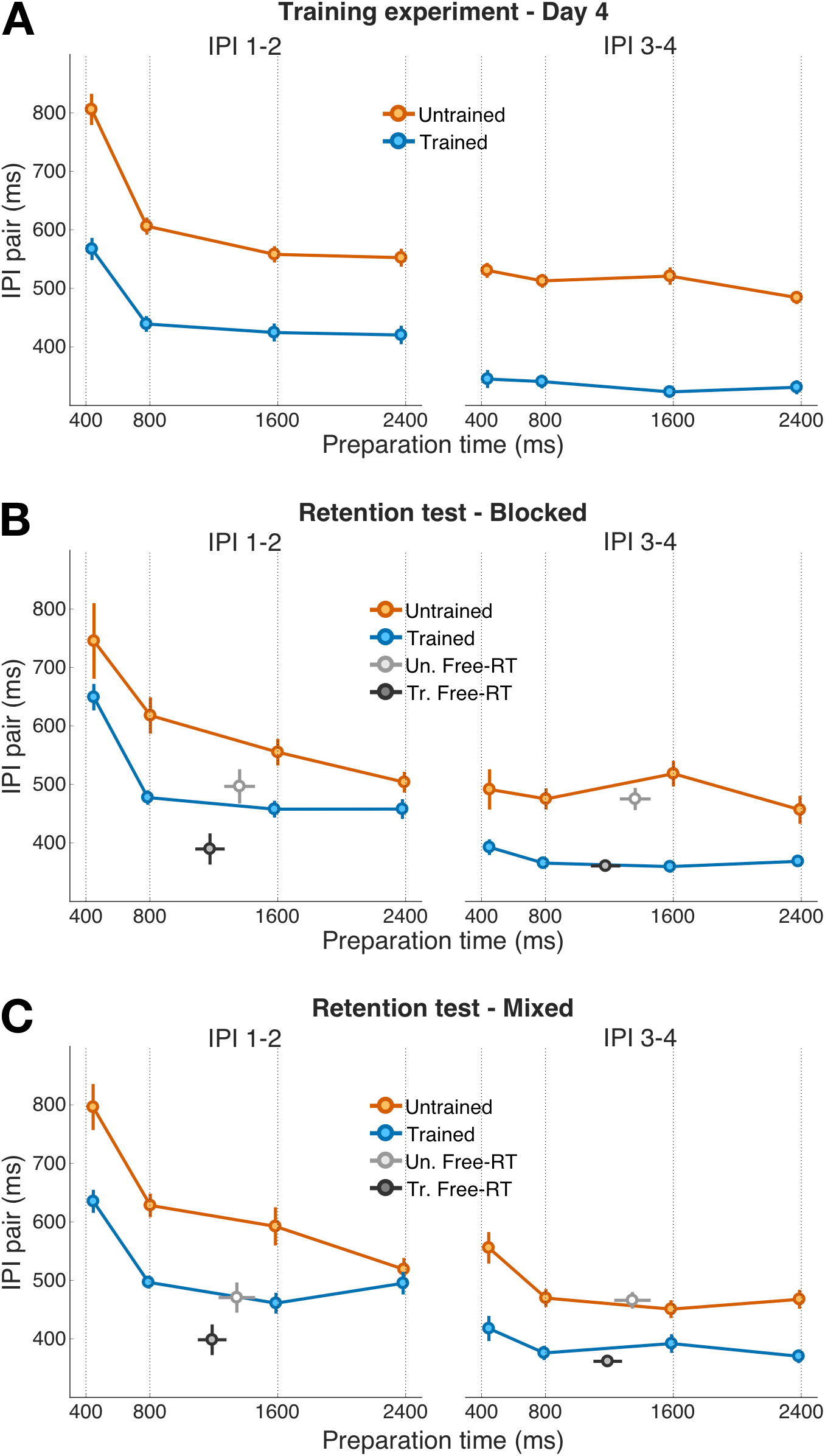
Improved online planning robustly underlies sequence-specific learning. Mean duration of IPI pairs (first 2 IPIs left, last 2 IPIs right) as a function of preparation time, separately for trained (blue) and untrained (orange) sequences. Other figure conventions are the same as in Figure 5A. **A.** Training experiment, Day 4, last 4 blocks (2 trained, 2 untrained). **B.** Retention test, Blocked blocks. For comparison, mean IPI pair durations for the Free-RT task are also shown in light gray and dark gray, for untrained and trained sequences respectively. **C.** Retention test, Mixed blocks.

In the training experiment (day 4, last 4 blocks: 2 trained, 2 untrained) we found that IPI 1-2 of trained sequences were significantly faster than those of the untrained sequences, for all preparation times (all preparation time comparisons: *t*_19_ > 5, *p* < 0.0001; Fig. 8A). However, in the retention test this was only true for the blocked paradigm (preparation time 2400 ms, IPI 1-2, *t*_14_ = 2.64, *p* = 0.019; Fig. 8B, left). In mixed blocks, we found no significant difference in mean ET between trained and untrained sequences (preparation time 2400ms, IPI 1-2, mean difference: 24 ± 25 ms; *t*_14_ = 0.95, *p* = 0.36; Fig. 8C, left). Overall, these mixed results point to a limited role of motor execution processes to sequence learning.

However, for IPI 3-4, we found a robust main effect of sequence type for the longest preparation time (Fig. 8, right), regardless of experiment (training: *F*_1,19_ = 91.21, *p* < 0.0001; retention: *F*_1,14_ = 65.51, *p* < 0.0001), block type (blocked: *F*_1,14_ = 49.98, *p* < 0.0001; mixed: *F*_1,14_ = 47.62, *p* < 0.0001), or paradigm (Forced-RT: *F*_1,14_ = 68.34, *p* < 0.0001; Free-RT: *t*_14_ = 6.32, *p* < 0.0001). Furthermore in the retention test a within-subject ANOVA on ET at 2400 ms (full preparation) revealed a significant interaction between IPI pair (1-2 vs. 3-4) and sequence type (trained vs. untrained; *F*_1,14_ = 5.09, *p* = 0.041), indicating that at 2400 ms there was a bigger sequence-type effect for IPI 3-4 than for IPI 1-2. Thus, when pre-planning is complete (IPI 1-2) the ET advantage for trained sequences was small; whereas when online planning played a bigger role (IPI 3-4) trained sequences were executed consistently faster than untrained ones. This demonstrates that 1) online planning improved as a consequence of sequence training, and that 2) improvements in online planning, rather than in motor execution or sequence pre-planning, underlie most of the sequence-specific execution advantage observed at the end of training.

## Discussion

Sensorimotor sequence learning manifests when individual action elements can be accurately executed in rapid succession (Beukema and Verstynen 2018; Diedrichsen and Kornysheva 2015; Verwey and Wright 2014). We combined the DSP task with a Forced-RT paradigm to uncover which elementary components supported such skilled motor performance, and how they change with learning. One element of skill is the selection and initiation of a single response (out of 5 possible alternatives), an ability that improves with practice (Haith et al. 2016; Hardwick et al. 2017). We also show that on average participants pre-plan the first 3 sequence elements, and can do so in less than 2 seconds. The speed of pre-planning improves with sequence-specific, but not sequence-general, learning. However, regardless of the amount of time allowed to prepare, the pre-planning capacity (i.e., how many actions can be pre-planned into the future) appears to be limited. Therefore, for sequences longer than 3 items, or when given little time to prepare, the remaining sequence elements have to be planned online during the execution of the beginning of the sequence. We show this process is faster for trained than untrained sequences. Overall, our results support the view that sequence-specific learning is explained by improvements in planning processes both before (i.e., pre-planning) and during sequence execution (i.e., online planning).

### Quicker single-item selection underlies sequence-general learning

Many studies show that behavioral training improves performance in the execution of repeating (i.e., trained) sequences. However, in the DSP task, there are also substantial performance improvements for the execution of novel (i.e., untrained) sequences (Fig. 3; Kornysheva and Diedrichsen, 2014; Waters-Metenier et al., 2014; Wiestler et al., 2014). One component of this sequence-general learning is the process of action selection: regardless of the specific sequence order, participants become better at mapping visual stimuli to motor responses for each individual finger press. Indeed, consistent with previous findings (Haith et al., 2016; Hardwick et al., 2017) we show that the speed and accuracy of selection (or S-R mapping) of individual items improved with practice. Thus, faster selection of individual items can account for sequence-general learning in the context of the SRT and DSP tasks. However, given that it is defined to work on one element at a time, this process cannot account for sequence-specific learning effects.

### Pre-planning of multiple items leads to faster sequence execution

In our task, all sequence elements are shown to the participant before the start of the execution. Therefore, other than improvements in single-item selection, performance benefits could also arise from an improved ability to pre-plan multiple sequential movements prior to execution (Magnuson et al. 2008; O’Shea and Shenoy 2016; Sheahan et al. 2016; Verwey 1994, 2001; Wong et al. 2015a). Indeed, given enough time before sequence initiation, pre-planning does not stop at the first element of the motor sequence (Verwey et al. 2010, 2014). Instead, participants tend to look ahead and pre-plan more than one sequence element at a time. By combining the DSP task with a Forced-RT paradigm, we obtained an indirect measure sequence planning (Haith et al. 2016): short preparation times resulted in slower execution for the first few IPIs, as compared to longer preparation times. From these results we infer that more motor planning has occurred during longer preparation times. Given that execution times seemed to reach an asymptote, it is likely that this process was fully completed by 2400 ms. However, even when provided with the longest preparation time, participants only prepared the first 3-4 elements of the sequence.

What does determine this upper limit of pre-planning capacity that we observed? Given the capacity of working memory (Miller 1956), it may be somewhat surprising that participants could not fully pre-plan a 5-item sequence, especially after a multi-day training period. It is well possible that this result reflects a general limitation of the motor system to only be able to pre-plan 3-4 elementary movements into the future. Another possibility is that this limit in pre-planning capacity was in fact dictated by the dual-task requirements of our forced-response paradigm (Ghez et al. 1997). This paradigm required participants to attend to two demands concurrently: to initiate a response in synchrony with the last in a series of tones, and to pre-plan the correct finger presses. Thus, it may be that under unconstrained conditions people can actually pre-plan a few more elements. Our Free-RT condition in the retention test shows that reaction times can be faster than long preparation times in the Forced-RT condition while still reaching comparable sequence execution speed (Fig. 7A-7C). However, a closer inspection of the IPI profiles (Fig. 7B-7D) suggests that the difference in execution speed between Free- and Forced-RT conditions is mostly driven by the first IPI (i.e., the first 2 presses). In fact, in the retention test we did not find a difference on the last 2 IPIs (of either trained or untrained sequences) between Free- and Forced-RT conditions. This indicates that 1) 2400 ms is sufficient to reach pre-planning capacity, and that 2) dual task requirements of the Forced-RT paradigm did not limit pre-planning capacity.

Results obtained from both invasive recordings in non-human primates and neuroimaging in humans under unconstrained conditions also seem to support the notion of a limited pre-planning capacity (Averbeck et al. 2002; Kornysheva et al. 2018). In accordance with the competitive queuing hypothesis (for a review, Rhodes et al., 2004), these studies show that 3-4 specific segments of sequential reaching movements are already pre-activated before movement onset, with the strength of representation for each segment reflecting the serial position of the planned element in the motor sequence.

The main implication of this idea is that, if the length of the sequence exceeds the pre-planning capacity, later sequence elements need to be selected and planned during the ongoing execution of earlier sequence elements. This is clearly evident from Fig. 6C, showing that the first two IPIs were significantly faster in the 2400 ms vs. 400 ms preparation time condition (an index of superior pre-planning), whereas the last two IPIs were not. Therefore sequence planning does not end at the onset of execution (Verwey 1996), but continues during execution as online planning.

### Sequence training improves pre-planning ability

We then investigated whether pre-planning, online-planning, and motor execution processes improved with sequence-specific learning. Our first novel result, consistent across training and retention experiments, was that pre-planning became faster for trained sequences (Fig. 6B). In other words, participants needed less time to reach a fully prepared state (i.e., to reach planning capacity). As a consequence, longer preparation times were only beneficial for the execution of unfamiliar sequences. The faster pre-planning was also evident in faster reaction times when movement initiation was not constrained in the retention experiment (i.e., Free-RT paradigm). Interestingly we found that, despite the overall gain in execution speed, there was no change in pre-planning speed for untrained sequences between day 1 and day 4 (Fig. 6B), suggesting that the ability to improve pre-planning is sequence-specific and depends on sequence familiarity (developed through training). Furthermore, our data indicated that the pre-planning capacity of 3-4 items was relatively fixed and could not be increased with training.

### Does sequence-specific learning affect motor execution skills?

In the training experiment we found that trained sequences were executed faster than untrained ones even when participants were provided with ample time to prepare the sequences (Fig. 6A). This effect was also observed for the first two IPIs, which participants could fully prepare (Fig. 8A), and for which online planning should play a minor role. These findings could therefore be taken as evidence that sequence-specific learning improves motor execution itself, beyond the benefit of sequence pre-planning.

What could explain such an improvement in motor execution? It should be noted that trained and untrained sequences were matched for probability of 1^st^ order transitions. That is, learning to execute specific transitions between any two fingers should have benefitted the production of trained and untrained sequences equally. A putative sequence-specific learning effect on the execution level, therefore must consist of learning specific transitions between 3 or more fingers.

While there was a clear execution advantage for the first two IPIs of trained sequences in the training experiment and the blocked retest, we did not replicate this finding in the mixed retest (Fig. 8C, left): the difference between IPI 1-2 of trained and untrained sequences for the longest preparation time (2400 ms) was not significant. Therefore, our results remain somewhat inconclusive in respect to sequence-specific improvements in motor execution. Although learning to execute specific 3-finger transitions may have played a role in speed improvements, the result may be more pronounced in blocked conditions (Fig. 7A-7C). This may suggest a role of increased motivation in blocks with only trained sequences (Wong et al. 2015b). Alternatively, the difference could indicate that sequence-specific execution skills are not as readily recalled under the uncertainty of a mixed block.

### Improved online planning drives sequence learning effects

Even though the sequence-specific advantage of trained over untrained sequences was not always evident on the first two IPIs, we found clear evidence for a strong and robust difference on subsequent IPIs. Indeed, we showed that the speed difference between sequence types was more pronounced for the end, as compared to the beginning, of the sequence. The most likely origin of this effect is that IPI 1-2 could be pre-planned in advance (and comparably well for either sequence type), whereas IPIs 3-4 required online planning, which became faster with training.

The sequence-specific improvement in online planning could have two explanations. First, as it happens for pre-planning, selection and planning processes may become more efficient when acting on a trained sequence. Alternatively, execution of well-learned sequences may require less central attentional resources, and instead rely more on the autonomous “motor processor” (Verwey 2001). The “cognitive processor” would therefore have more resources to look ahead and prepare the next responses, i.e., the possible interference between planning and execution processes would thus be reduced. Either way, our results indicate that sequence-specific learning likely occurs through improvements at the very interface between cognitive (selection) and motor (execution) processes (Diedrichsen and Kornysheva 2015).

### Conclusions

The combination of experimental approaches used here allowed us to disentangle different components of skill learning in a sequence production task. We showed that performance improvements cannot be fully explained by either faster single-item selection, nor improved motor execution. Instead, much of the sequence-specific advantage can be attributed to an enhanced ability to select and plan multiple sequence elements into the future, evident both before sequence production starts, as well as during sequence execution. Whether these findings generalize to other types of actions (e.g., reaching) remains to be seen. If they do, it would open up the possibility to investigate the neural mechanisms of online planning in animal models. For such experiments, however, it will be critical to ensure that the number of sequence elements is higher than the pre-planning capacity, such that online planning and motor execution processes can be dissociated. Overall, online motor planning constitutes a central mechanism at the interface between the cognitive and the motor system that allows the brain to deal with a continuous stream of stimuli and motor demands.

Author contributions
G.A. and J.D. designed research; G.A. performed research; G.A. analyzed data; G.A. and J.D. wrote the paper.

## Acknowledgements

This work was supported by a James S. McDonnell Foundation Scholar award, an NSERC Discovery Grant (RGPIN-2016-04890), and the Canada First Research Excellence Fund (BrainsCAN). The authors wish to thank Eva Berlot and Spencer Arbuckle for comments on the manuscript.

## References

Abrahamse EL, Ruitenberg MFL, de Kleine E, Verwey WB. Control of automated behavior: insights from the discrete sequence production task. Front Hum Neurosci 7 1–16, 2013.

Averbeck BB, Chafee M V., Crowe DA, Georgopoulos AP. Parallel processing of serial movements in prefrontal cortex. Proc Natl Acad Sci 99: 13172–13177, 2002.

Beukema P, Verstynen T. Predicting and binding: interacting algorithms supporting the consolidation of sequential motor skills. Curr Opin Behav Sci 20: 98–103, 2018.

Diedrichsen J, Kornysheva K. Motor skill learning between selection and execution. Trends Cogn Sci 19: 227–233, 2015.

Doyon J, Gabitov E, Vahdat S, Lungu O, Boutin A. Current issues related to motor sequence learning in humans. Curr Opin Behav Sci 20: 89–97, 2018.

Ghez C, Favilla M, Ghilardi MF, Gordon J, Bermejo R, Pullman S. Discrete and continuous planning of hand movements and isometric force trajectories. Exp Brain Res 115: 217–233, 1997.

Haith AM, Pakpoor J, Krakauer JW. Independence of Movement Preparation and Movement Initiation. J Neurosci 36: 3007–3015, 2016.

Hardwick RM, Forrence AD, Krakauer JW, Haith AM. Skill Acquisition and Habit Formation as Distinct Effects of Practice. bioRxiv 1–35, 2017.

Kornysheva K, Bush D, Meyer S, Sadnicka A, Barnes G, Burgess N. Neural competitive queuing of ordinal structure underlies skilled sequential action. bioRxiv, 2018.

Kornysheva K, Diedrichsen J. Human premotor areas parse sequences into their spatial and temporal features. Elife 3: e03043, 2014.

Magnuson CE, Robin DA, Wright DL. Motor programming when sequencing multiple elements of the same duration. J Mot Behav 40: 532–544, 2008.

Miller GA. The magical number seven, plus or minus two: some limits on our capacity for processing information. Psychol Rev 63: 81–97, 1956.

O’Shea DJ, Shenoy K V. The Importance of Planning in Motor Learning. Neuron 92: 669–671, 2016.

Rhodes BJ, Bullock D, Verwey WB, Averbeck BB, Page MPA. Learning and production of movement sequences: Behavioral, neurophysiological, and modeling perspectives. Hum Mov Sci 23: 699–746, 2004.

Sheahan HR, Franklin DW, Wolpert DM. Motor Planning, Not Execution, Separates Motor Memories. Neuron 92: 773–779, 2016.

Shmuelof L, Krakauer JW, Mazzoni P. How is a motor skill learned? Change and invariance at the levels of task success and trajectory control. J Neurophysiol 108: 578–594, 2012.

Verwey WB. Evidence for the development of concurrent processing in a sequential keypressing task. Acta Psychol (Amst), 1994. doi:10.1016/0001-6918(94)90038-8.

Verwey WB. Buffer Loading and Chunking in Sequential Keypressing. J Exp Psychol Hum Percept Perform, 1996. doi:10.1037/0096-1523.22.3.544.

Verwey WB. Concatenating familiar movement sequences: The versatile cognitive processor. Acta Psychol (Amst) 106: 69–95, 2001.

Verwey WB, Abrahamse EL, de Kleine E. Cognitive processing in new and practiced discrete keying sequences. Front Psychol 1: 1–13, 2010.

Verwey WB, Shea CH, Wright DL. A cognitive framework for explaining serial processing and sequence execution strategies. Psychon Bull Rev 22: 54–77, 2014.

Verwey WB, Wright DL. Learning a keying sequence you never executed: Evidence for independent associative and motor chunk learning. Acta Psychol (Amst) 151: 24–31, 2014.

Waters-Metenier S, Husain M, Wiestler T, Diedrichsen J. Bihemispheric Transcranial Direct Current Stimulation Enhances Effector-Independent Representations of Motor Synergy and Sequence Learning. J Neurosci 34: 1037–1050, 2014.

Wiestler T, Diedrichsen J. Skill learning strengthens cortical representations of motor sequences. Elife 2: e00801, 2013.

Wiestler T, Waters-Metenier S, Diedrichsen J. Effector-Independent Motor Sequence Representations Exist in Extrinsic and Intrinsic Reference Frames. J Neurosci 34: 5054–5064, 2014.

Wong AL, Haith AM, Krakauer JW. Motor Planning. Neurosci 21: 385–398, 2015a.

Wong AL, Lindquist MA, Haith AM, Krakauer JW. Explicit knowledge enhances motor vigor and performance: motivation versus practice in sequence tasks. J Neurophysiol 114: 219–32, 2015b.

Yokoi A, Arbuckle SA, Diedrichsen J. The role of human primary motor cortex in the production of skilled finger sequences. J Neurosci 38: 2798–17, 2018.

Yokoi A, Bai W, Diedrichsen J. Restricted transfer of learning between unimanual and bimanual finger sequences. J Neurophysiol 117: 1043–1051, 2017.

